# DeepMapper: Attention-Based AutoEncoder for System Identification in Wound Healing and Stage Prediction

**DOI:** 10.1101/2024.12.17.628977

**Authors:** Fan Lu, Ksenia Zlobina, Sebastian Osorio, Hsin-ya Yang, Alexandra Nava, Michelle D. Bagood, Marco Rolandi, Roslyn Rivkah Isseroff, Marcella Gomez

## Abstract

Advancements in bioelectronic sensors and actuators have paved the way for real-time monitoring and control of wound healing progression. Real-time monitoring allows for precise adjustment in treatment strategies that align with an individual’s unique biological response. However, due to the complexities of human-drug interactions and a lack of predictive models it is challenging to determine just how one should adjust drug dosage to achieve the desired biological response. This work proposes an adaptive closed-loop control framework that integrates deep learning, optimal control, and reinforcement learning to update treatment strategies in real-time with the goal of accelerating wound closure. The proposed approach eliminates the need for mathematical modeling of complex nonlinear wound healing dynamics. We demonstrate the convergence of the controller via an *in silico* experimental setup, where the proposed approach successfully accelerates the wound healing process by 17.71%. Finally, we share the experimental setup and results of an *in vivo* implementation to highlight the translational potential of our work. Our data-driven model estimates a 40% acceleration in wound closure.

## Introduction

Wound healing is a complex, nonlinear biological process that involves multiple stages and relies on various cellular and biochemical interactions. Accurately predicting the healing stages of wounds is crucial for effective treatments, as it allows clinicians to intervene appropriately based on the current state of healing. However, due to the inherent nonlinear dynamics of wound healing, traditional linear methods often fall short in capturing the nuanced dependencies and temporal interactions within the healing process.

Wound healing progresses through four main stages: hemostasis, inflammation, proliferation, and maturation (1). While these stages can be modeled as linear dynamics (2), it is hard to accurately determine the wound’s state at any given time. One key challenge is the lack of labels for wound stages, as labeling can be highly subjective, with different clinicians interpreting the wound’s stage differently. Additionally, such a labeling process can be time-consuming and expensive, making it hard to build a supervised machine learning model for wound stage prediction.

Recently, some unsupervised learning approaches have been proposed to address the scarcity of labeled data (3). While these approaches show promise, they struggle with generalization and lacks consideration of the temporal relationships between wound stages. Similarly, (4) introduced a model that incorporated temporal data, but it was only validated on a specific species (mice), limiting its broader application and generalization. These limitations highlight the need for more robust models that can capture the temporal and feature dependencies of wound healing across different datasets and species.

This motivated the design of a new deep neural network-based algorithm, DeepMapper, which combines the strengths of AutoEncoders and attention mechanisms to learn a robust mapping from unknown nonlinear wound healing dynamics to their linear representations, similar to the goal of the Koopman Operator (5). DeepMapper seeks to capture the temporal and feature dependencies in wound healing dynamics, allowing for a linear representation that facilitates understanding, tracking, and predicting the healing stages. By leveraging both AutoEncoder structures for efficient feature encoding and transformer-based attention mechanisms for enhanced sequence modeling (6), DeepMapper aims to create interpretable, scalable representations that advance the modeling of wound healing processes.

Learning linear representations of nonlinear systems is essential as nonlinear dynamics are pervasive in real-world applications, from biological processes to complex physical systems. Unlike linear systems, which have well-established methods for design, control, analysis, and optimization (7, 8), nonlinear systems present challenges due to their complexity and lack of a general, scalable framework for analysis (9–11). The ability to find a reliable linear representation can also enable us to leverage robust linear control techniques while managing the complexity inherent in nonlinear behaviors.

The challenge lies in accurately mapping these nonlinear dynamics to a linear space. The Koopman operator theory offers a powerful approach by representing a nonlinear system as an infinite-dimensional linear system, allowing us to apply linear analysis to nonlinear dynamics. Introduced by Koopman (5) and further developed in (12, 13), the Koopman operator effectively lifts nonlinear behaviors to a higher-dimensional linear space. However, practical applications are limited by the difficulty of optimizing over functional spaces, making direct Koopman representations computationally intractable. In this work, we harness deep learning to approximate the Koopman operator, constructing flexible representations that enforce parsimony and interpretability, enabling practical applications.

Neural networks (NNs) have the capacity of representing any continuous function to an arbitrary degree of accuracy, with sufficient hidden units and a linear output layer, according to the universal approximation theorem. This theoretical foundation underpins the use of neural networks in modeling complex systems, including the approximation of the Koopman operator. Building on the Koopman operator theory, recent research has extended its application to nonlinear systems by using AutoEncoders and other NN-based architectures. These methods employ AutoEncoders to encode complex behaviors into a latent space where linear representations of the underlying nonlinear system can be found (2, 9, 14–16). For multi-feature time series data, however, these methods overlook essential temporal and inter-feature relationships, reducing their effectiveness in capturing the dependencies that drive system evolution.

Recurrent neural networks RNNs—such as long short-term memory (LSTM) networks (17) and gated recurrent units (GRU) (18)—were commonly used for sequence modeling due to their ability to capture temporal dependencies. However, RNNs process data point-by-point across time steps, accumulating hidden states that depend on previous inputs. This sequential processing approach makes RNNs challenging to parallelize, limiting their scalability and slowing down performance, especially for long sequences.

The transformer model, an NN architecture based on attention mechanisms, offers a solution to these limitations by forgoing sequential processing in favor of parallelization. (6). Unlike recurrent neural networks (RNNs), transformers eliminate the sequential processing constraint by replacing recurrent structures with an attention mechanism, allowing for parallel computation across time steps. While time hierarchy is removed, transformers employ positional embeddings to preserve the order of sequential data points. This approach not only supports parallelization but also provides an efficient mechanism for modeling multi-feature time series, making transformers a powerful tool for understanding and predicting complex temporal relationships within nonlinear systems.

The main contributions of this paper include:

- We introduce DeepMapper, an attention-based autoencoder designed to learn a linear representation of nonlinear wound healing dynamics, through which wound stage will be accurately predicted.
- We demonstrate improved data efficiency in crossspecies learning, enabling the model to generalize across different biological contexts with reduced data requirements.
- Experimental results show that our method accurately forecasts the time required for full wound closure based on the learned linear model. The learned linear model can be used as a guide for doctors in the treatment planning of patients.

### Approach

While the visual dynamics of wound healing may appear non-linear, the overall process of transitioning through the four stages: hemostasis, inflammation, proliferation, and maturation (19–21), can be approximated by linear dynamics under certain conditions (2). If a linear representation could be derived to capture this nonlinear process, it could significantly enhance the predictability and interpretability of wound healing stages. Such a linear model would offer a powerful tool for simplifying the complex dynamics of wound healing, providing clinicians and researchers with a clearer and more actionable framework for analyzing wound progression and optimizing treatment strategies.

We propose a deep learning framework called DeepMapper to learn a mapping from a nonlinear wound healing dynamics to its linear representation. We show that the learned linear model can be used for accurate wound stage prediction. The learning framework is schematized in Fig. 1.

**Fig. 1.**
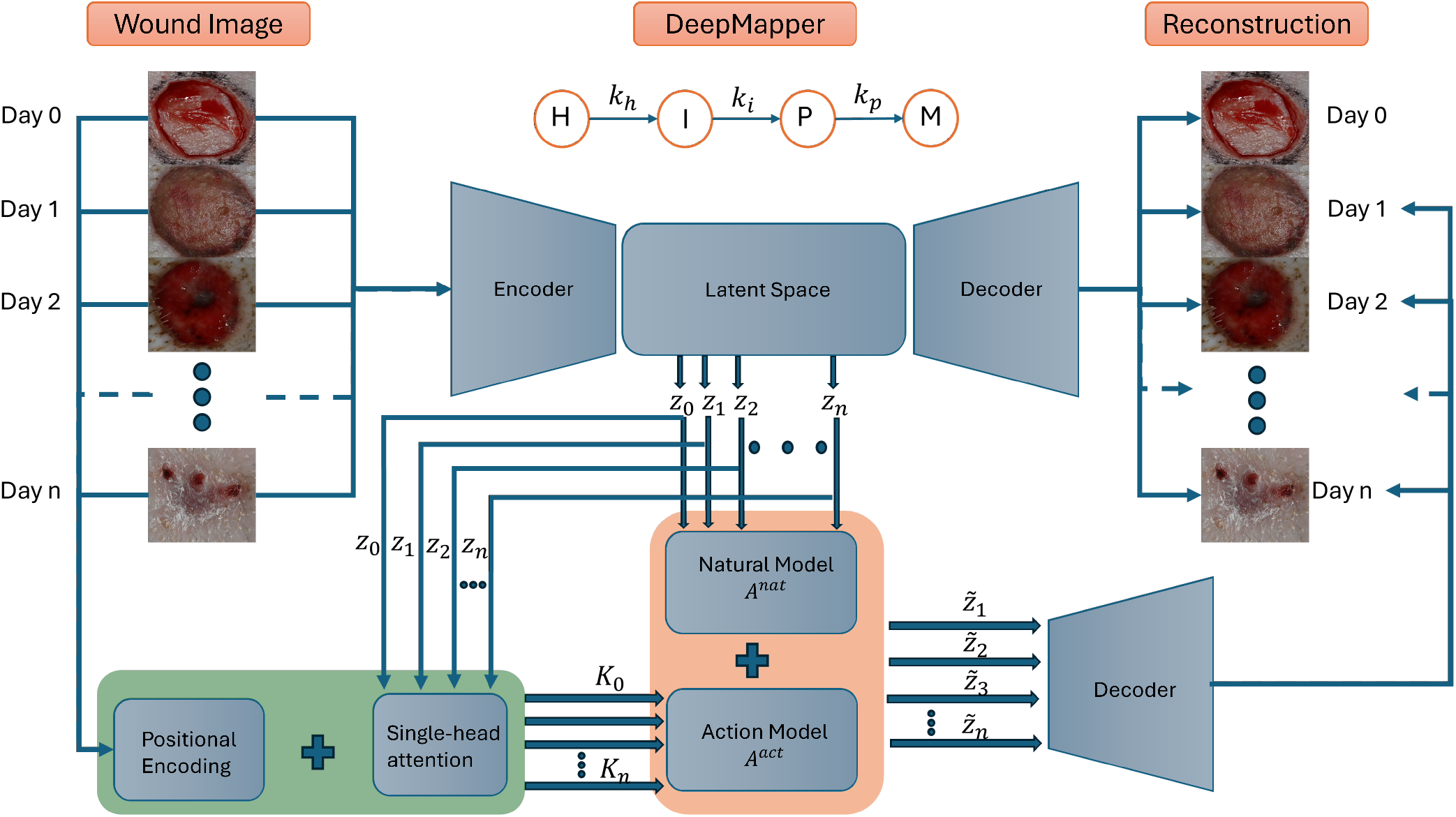
DeepMapper framework. The encoder of the DeepMapper takes in a sequence of wound images, mapping them to a sequence of variables *z* in the latent space. These variables are then passed into the Natural Model to learn the matrix *A*^act^ in the linear model. As *A*^act^ can be noisy, single-head attention model is then used to de-noise *A*^act^. The decoder of the DeepMapper then generates the corresponding wound images through variables in the latent space.

#### ODE Model Design

We assume that there are four variables in the linear representation, and each corresponds to the probability of each stage of wound healing: hemostasis *H*, inflammation *I*, proliferation *P*, and maturation *M*. As time goes on, the probability of each stage changes: the wound is initially in the hemostasis stage with probability one and then transitions through each consecutive stage:

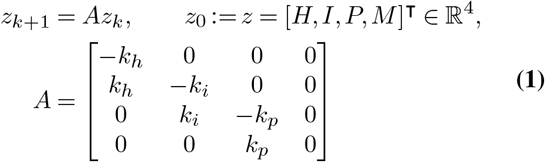

where *k*_*h*_, *k*_*i*_, *k*_*p*_ ∈ ℝ are the values that control the velocities of transitions from homeostasis to inflammation, from inflammation to proliferation, and from proliferation to maturation, respectively.

#### DeepMapper: linearization of nonlinear wound-healing dynamics

In practice, the wound state, denoted as *x* ∈ ℝ^*n*^, may be comprised of more than 4 variables, and often evolves nonlinearly. One example is a photo of the wound taken at time *k*, representing the wound state *x*_*k*_. Suppose the system follow a discrete-time dynamical system:

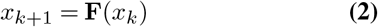

where *x* ∈ ℝ^*n*^ is the state of the system and **F** represents the dynamics that map the state of the system forward in time. Discrete-time dynamics often describes a continuous-time system that is sampled discretely in time, so that 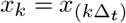 with sampling time Δ_*t*_. The dynamics in **F** are generally non-linear, and the state *x* may be high dimensional, although we typically assume that the dynamics evolve on a low-dimensional attractor governed by persistent coherent structures in the state space. Note that **F** is often unknown and only measurements of the dynamics are available.

The dominant geometric perspective of dynamical systems concerns the organization of trajectories of Eq. (2), including fixed points, periodic orbits, and attractors. Formulating the dynamics as a system of differential equations in *x* often admits compact and efficient representations f or many natural systems; for example, Newton’s second law is naturally expressed by Eq. (2). However, the solution to these dynamics may be arbitrarily complicated, and possibly even irrepresentable, except for special classes of systems. Linear dynamics, where the map **F** is a matrix that advances the state *x*, are among the few systems that admit a universal solution, in terms of the eigenvalues and eigenvectors of the matrix **F**, also known as the spectral expansion.

#### Koopman operator theory

In 1931, B.O. Koopman provided an alternative description of dynamical systems in terms of the evolution of functions in the Reproducing Hilbert Kernel space (RKHS) of possible measurements *z* = *h*(*x*) of the state (5). The so-called Koopman operator, 𝒦, that advances measurement functions is an infinite-dimensional linear operator:

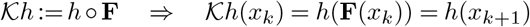

Koopman analysis has gained significant attention recently, largely due to the pioneering work of (12, 22), driven by the increasing availability of measurement data and the lack of known equations governing many complex systems. By representing nonlinear dynamics within a linear framework through the Koopman operator, this approach holds the potential to facilitate advanced nonlinear prediction, estimation, and control using the well-established theory for linear systems. However, obtaining finite-dimensional approximations of the infinite-dimensional Koopman operator remains a significant challenge in practical applications, which strongly motivates the adoption of deep learning methods, given their universal function approximation capabilities.

#### Deep learning to identify Koopman operators

The primary goal of this work is to leverage deep neural networks to approximate a function *h* : ℝ^*n*^ → ℝ^*d*^ that serves a similar role to the Koopman operator, mapping nonlinear states in Eq. (2) to linear ones in Eq. (1). Our approach is driven by the need for parsimonious representations that are efficient, resist overfitting, and provide compact, interpretable descriptions of the system’s dynamics in intrinsic coordinates. Unlike previous deep learning-based Koopman methods, our network architecture is specifically designed to capture temporal relationships in the data while updating a prior model of a predesigned linear system, ensuring domain-specific interpretability, particularly in wound healing.

Our core network architecture is shown in Fig. 1. The objective of this network is to identify a mapping function *h* : ℝ^*n*^ → ℝ^*d*^, along with a dynamical system *z*_*k*+1_ = *Az*_*k*_. There are three high-level requirements for the network, corresponding to three types of loss function used in training.

#### Intrinsic coordinates that are useful for reconstruction

We seek to identify a few intrinsic coordinates *z* = *h*(*x*) where the dynamics evolve, along with the inverse *x* = *h*^*−*1^(*z*) so that the state *x* may be recovered. This is achieved using an auto-encoder (see Fig. 2), where *h* is the encoder and *h*^*−*1^ is the decoder. The dimension *d* of the auto-encoder latent space is a byperparameter of the network, and this choice may be guided by knowledge of the system, e.g., *d* = 4 in wound healing Eq. (1). Reconstruction accuracy of the auto-encoder is achieved via minimizing the loss:

**Fig. 2.**
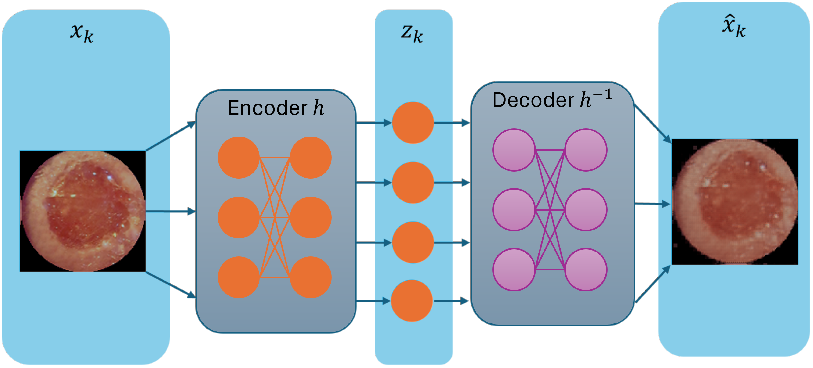
Reconstruction of DeepMapper: DeepMapper contains a encoder that maps the nonlinear state *x*_*k*_ into linear state *z*_*k*_. The nonlinear state is reconstructed by the decoder of the DeepMapper.

**Fig. 3.**
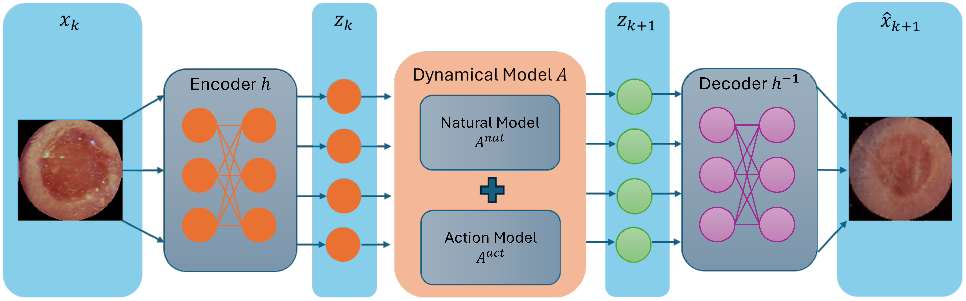
Reconstruction of DeepMapper with one step looking ahead. The linear state combined with the transition matrix *A* gives the next linear state *z*_*k*+1_ following Eq. (1). The next nonlinear state is reconstructed by the decoder of the DeepMapper.

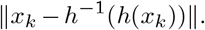

with ∥·∥ the mean-squared error, averaging over dimension then number of examples.

To enhance the resolution of the decoder’s output, high-resolution features from the contracting path of encoder are fused with the upsampled output from the decoder, following the approach used in the U-Net architecture (23).

#### Linear dynamics

To discover the mapping function *h*, we learn the transition matrix *A* in the linear dynamics Eq. (1) on the intrinsic coordinates, i.e., *z*_*k*+1_ = *Az*_*k*_. Linear dynamics are achieved via minimizing the following loss:

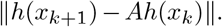

More generally, we enforce linear prediction over *m* time steps with the loss:

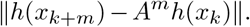

#### Future state prediction

Finally, the intrinsic coordinates must enable future state prediction. Specifically, we identify linear dynamics in the matrix *A*. This correspons to minizing the loss:

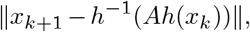

and more generally,

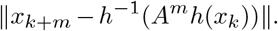

#### Attention Mechanism for learning linear models

In dynamcial systems where the state evolves over time, temporal attention mechanism allows the model to assign different importance to previous states. This helps in identifying systems that exhibit time-varying dynamics or where certain past events have a more significant influence on the present state than others (6).

The linear representation of the nonlinear wound healing dynamics described in Eq. (1) requires us to learn *k*_*h*_, *k*_*i*_, and *k*_*p*_ through a time series of wound images. Typically, these three parameters are proportional to the difference between two consecutive linear states *z*_*k*+1_ and *z*_*k*_ and can be estimated via Euler approximation. However, due to image noise and slow nature of wound healing (a typical wound takes around 14 days to close), such estimation is prone to significant inaccuracies. Instead, the aggregation of the previous several states will result in substantial changes of the wound states, both in linear and nonlinear dynamics. This motivates the use of an attention mechanism to capture the temporal relationships of each wound image, and thus gives a better prediction on the matrix *A*.

In the proposed approach shown in Fig. 1, instead of learning the *A* matrix directly through the attention model, we split it into two parts: *A*^nat^, which is learned directly from the latent space, and *A*^act^, which learns from the attention model to determine the correction needed to capture the observed dynamics:

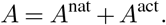

This approach effectively reduces noise during learning. In the experiment, we found that a single-head attention will give relatively good results without the need of a multi-head attention as in (6). The structure is shown in Fig. 4.

**Fig. 4.**
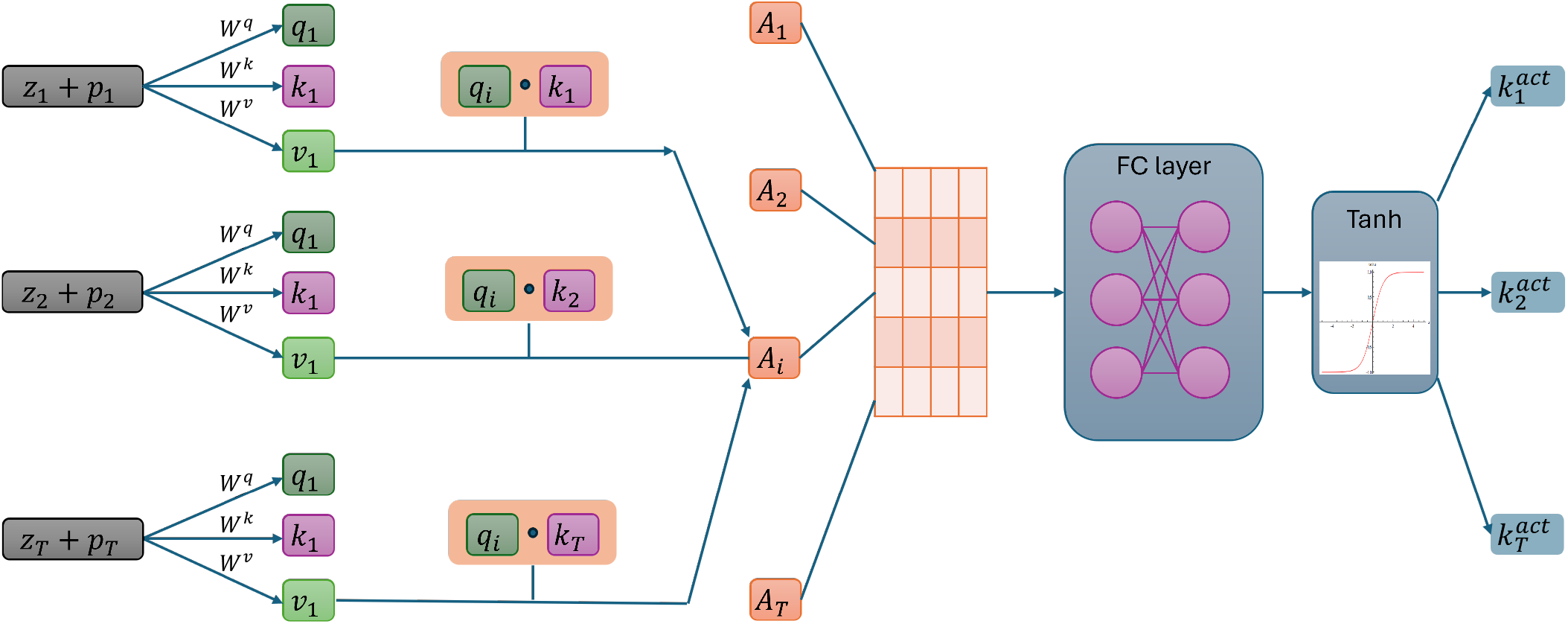
Single-head attention architecture. Three matrices with learnable weights, *W*^*Q*^, *W*^*K*^, and *W*^*V*^, are constructed to learn the temporal relationships of a time series of data. *x · y* represents the dot product of two vectors.

In order for the model to make use of the order of the sequence, we must inject some information about the relative or absolute temporal position of each image in the sequence. To this end, we add positional encodings to the input embeddings at the bottoms of the encoder and decoder stacks. The positional encodings have the same dimension *d* as the embeddings, so that the two can be summed. There are many choices of positional encoding, and we used the same as in (6):

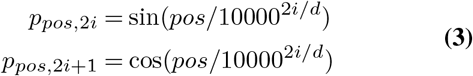

where *pos* is the position and *i* is the dimension. That is, each dimension of the positional encoding corresponds to a sinusoid. The wavelengths form a gemetric progression from 2*π* to 10000 · 2*π*. We chose this function because we hypothesized it would allow the model to easily learn to attend by relative positions, since for any fixed offset *k, p*_*pos*+*k*_ can be represented as linear function of *PE*_*pos*_.

We construct a query matrix *W*^*Q*^ ∈ ℝ^*N ×d*^ and a key matrix *W*^*K*^ ∈ ℝ^*N ×d*^, each with learnable weights from a deep neural network.

For each of the linear states *z*_*k*_ with 1 ≤*k* ≤ *T*, we compute *q*_*k*_ = *W*^*Q*^*z*_*k*_ to represent the query on how much the state *z*_*k*_ will affect other states. The product of *K* and *z*_*k*_ yields *a*_*k*_ = *Kz*_*k*_, which serves as the answer to the query. Then we apply the dot product and a Softmax function over the dot product to measure how close the answer to the query, and because the future state can only be affected by previous states, we apply a mask so that when *i < j, d*(*i, j*) = 0

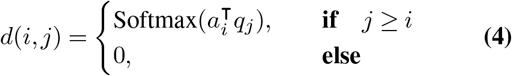

with 1 ≤*i, j* ≤ *N*.

We then define a final matrix *W*^*V*^, whose elements are from weights in neural networks. The value *v*_*t*_ = *W*^*V*^ *z*_*t*_ indicates the significance of each linear state to the others. Combining with *d*(*i, j*) gives the final attention score:

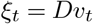

with *D* ∈ ℝ^*N ×N*^ the matrix containing all the attention score. This attended score will be passed into several linear layer to predict the three rates of transitions 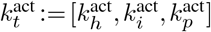, which are used to construct the matrix *A*^act^.

### Datasets and Experiment Results

We used two datasets to evaluate the performance of the proposed approach. One is the mouse wound dataset, and the other one is the porcine wound dataset. These datasets are split into training, validation, and testing datasets. Models are trained on training set and compared on the validation dataset, which is also used for early stopping to prevent over fitting. We report accuracy on the test set.

#### Mouse Dataset

The experiment for creating the mouse dataset used in this work is described in (24–26), from which we generate circular wound-only crops. The dataset includes wound images from eight mice, consisting of four mice from one cohort and four mice from a second distinct cohort, each imaged daily over a span of 16 days. Each individual had a wound inflicted on the left side and another on the right side. The resulting dataset is 256 images (8 mice *×* 2 wounds *×* 16 days).We wish to investigate whether our model can pick up on any subtle differences between the two cohorts and thus, demonstrate our model is not overfitting.

It is important to note that this dataset was not originally captured with computer vision applications in mind. Therefore, we had to address several challenges, including blur, occlusion, and illumination noise. To mitigate these issues during training, we applied various data augmentation techniques, such as random rotations and the addition of Gaussian noise. The dataset was divided into three subsets: 208 images for training, 16 for validation, and 32 for testing. The testing dataset includes two wound trajectories: T1 and T2. T1 represents a faster healing trajectory, while T2 demonstrates slower healing. T1 and T2 are selected such that they are not from the same cohort. This selection was based on the observed disparity in wound healing rates as illustrated in Fig. 5. This distinction is crucial for evaluating the effectiveness of our proposed method.

**Fig. 5.**
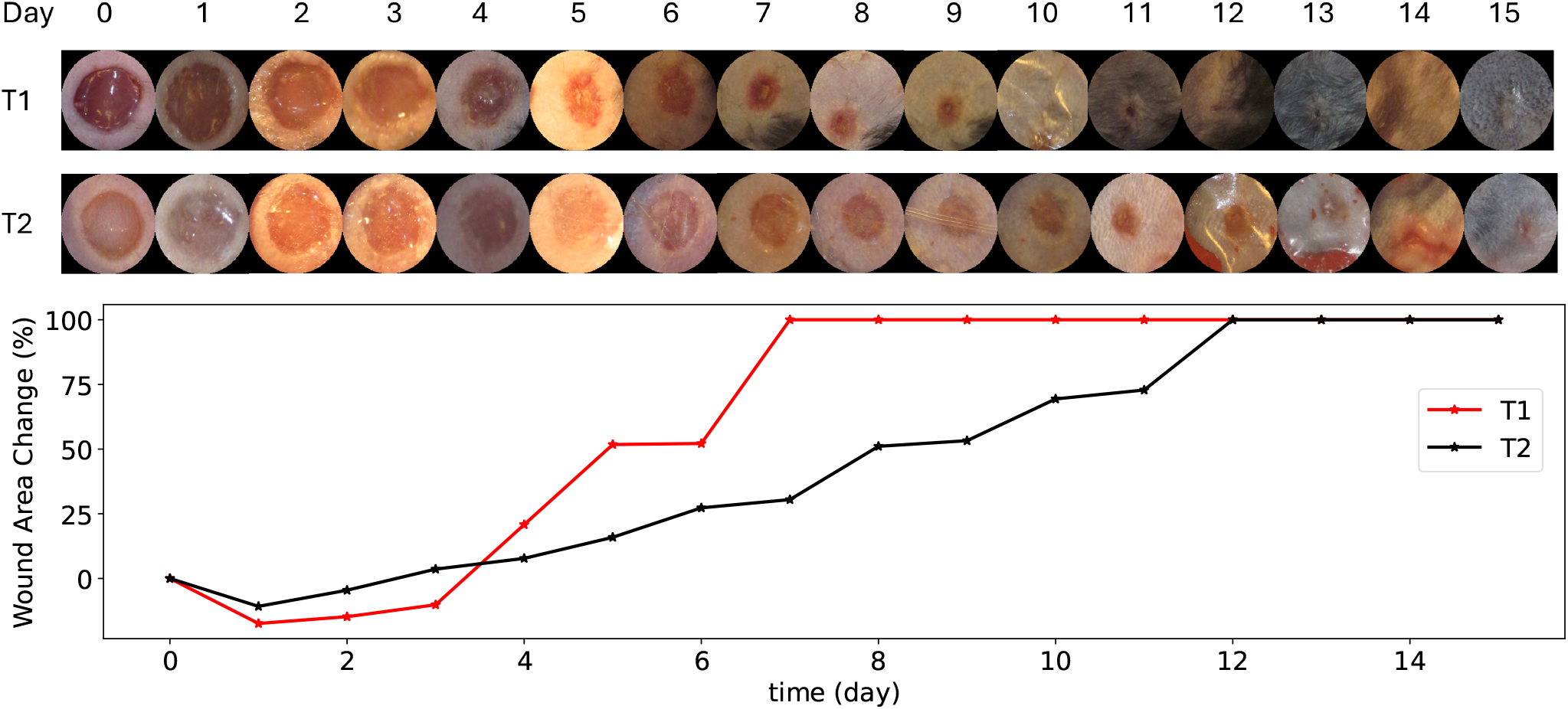
Two trajectories of testing wound images. T1: No visible wound remains after day 10. T2: The wound region persists beyond day 10, as evident from both the images and the wound area change plot.

#### Porcine Wound Dataset

The porcine wound dataset consists of wound images collected from six domestic pigs (Yorkshire-mix breed, female, 45-50 kg). In each pig, twelve full-thickness excisional wounds, each 2 cm in diameter, were created bilaterally—six on each side of the dorsum. No treatment was administered beyond basic wound dressing, allowing for typical, natural wound closure throughout the study (27).

During the postoperative period, wound images were captured using a digital single-lens reflex (DSLR) camera on Days 0, 1, 2, 3, 4, 5, 6, 7, 9, 11, 13, 15, 16, 19, and 21, with Day 0 marking the day of excision. For each observation, a 5-shot burst sequence was taken using the DSLR camera’s burst capture mode. To complement the imaging data, wound biopsies were performed at various stages of the experiment, limiting the number of wounds available for photography throughout the 21-day period. Consequently, only six wounds were consistently documented on Days 0, 1, 3, 5, 7, 11, 15, and 21, resulting in a total of 240 images captured using the burst-shot functionality of the DSLR camera.

The DSLR camera images also include a ruler, which can be used to estimate wound size—a critical factor for modeling wound healing progression. One example is shown in Fig. 7. We believe that incorporating this information is valuable for developing accurate linear models of wound healing. However, to simplify the experiments, we utilized images that only contain wound regions detected by the U-net model (23). This allowed us to focus solely on the wound area, avoiding the complexities of size estimation while ensuring consistency in the dataset used for analysis.

#### Experimental Results

For each experiment, We conduct 100 independent runs of training, with parameters in the neural networks randomly initialized via the Kaiming uniform method (28, 29). To get a better estimation of these values, we conduct Polyak–Ruppert averaging (30):

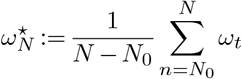

where *N* denotes the total number of updates in the parameters, and the interval [0, *N*_0_] with *N*_0_ *< N* is known as the *burn-in* period; estimates from this period are abandoned.

In order to show the robustness of our approach, we applied two training schemes: 1. DeepMapper trained from scratch using porcine wound dataset. 2. DeepMapper trained using the mouse wound dataset and fine-tuned on the porcine wound dataset. The objective of these training schemes was to evaluate DeepMapper’s ability to generalize across different species and to demonstrate the potential for knowledge transfer between species, thereby improving data efficiency. The results, presented in Fig. 8, indicate that fine-tuning led to a more stable learning process, with significantly faster convergence. This outcome underscores the model’s robustness in transferring knowledge across species and highlights its capacity for efficient data utilization.

The learned linear model parameters, when tested on the mouse dataset, are shown in Fig. 6. Using these parameters, we derived the evolution of the four wound stages over time, following the dynamics described in Eq. (1), as shown in Fig. 6. As can be seen from the lower-right plot of Fig. 6, the probability of maturation reaches 100% much more quickly in T1 mouse images compared to T2 mouse images, validating the accuracy of the learned parameters and aligning with the observed faster wound area shrinkage in T1 compared to T2. The learned linear model parameters, when tested on the porcine wound dataset, are shown in Fig. 9. Using these parameters, we derived the evolution of the four wound stages over time, following the dynamics described in Eq. (1), as shown in Fig. 9. As can be seen in Fig. 9, the probability of maturation for wound *I* converges at 21, closely matching the actual wound closure time.

**Fig. 6.**
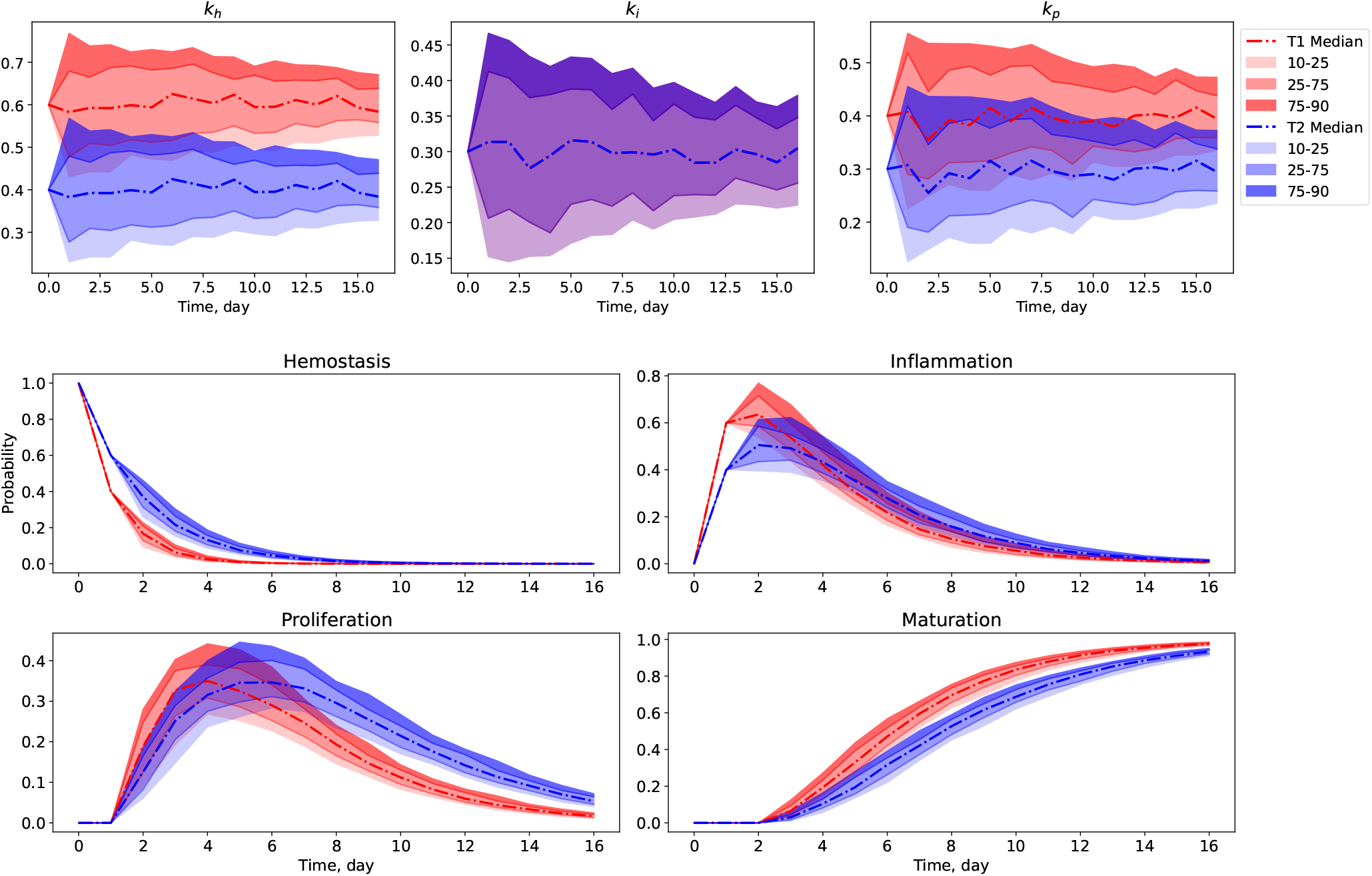
Experimental results of DeepMapper on Mouse dataset. Values of *k*_*h*_, *k*_*i*_, *k*_*p*_ and mouse wound stage prediction across time during testing, shown by percentile.

**Fig. 7.**
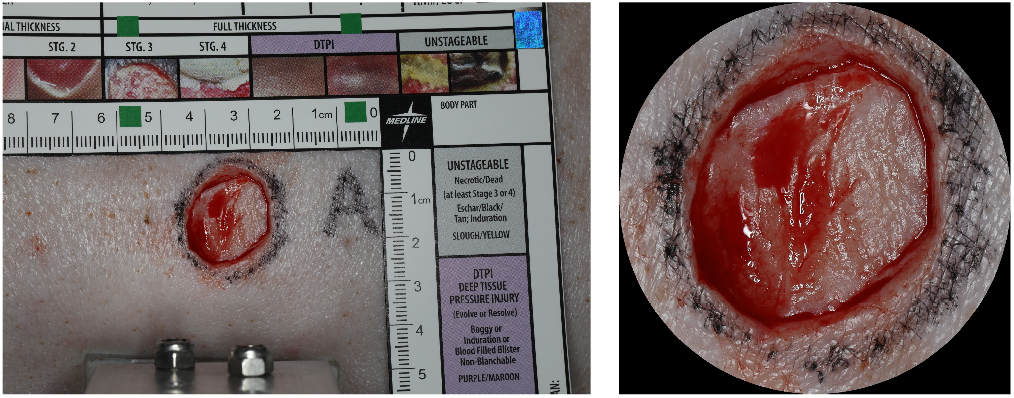
Sample image from porcine wound dataset: shown on the left is the original DSLR image; shown on the right is the image that only contains the wound region.

**Fig. 8.**
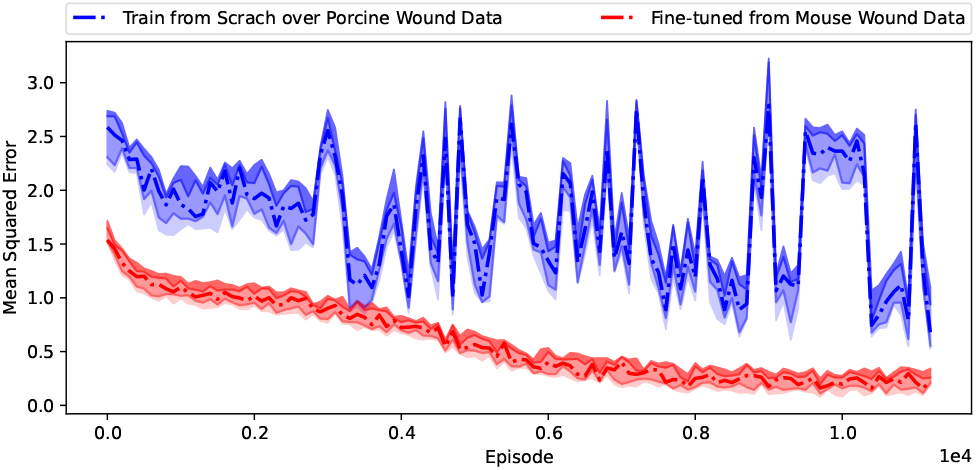
Learning curves of DeepMapper. Two training strategies were employed: (1) A randomized DeepMapper model directly trained over the porcine wound data. The learning curves are shown in the blue curves. (2) A pre-trained model over mouse wound dataset and fine-tuned over the porcine wound data. The learning curves are shown in the red curves.

**Fig. 9.**
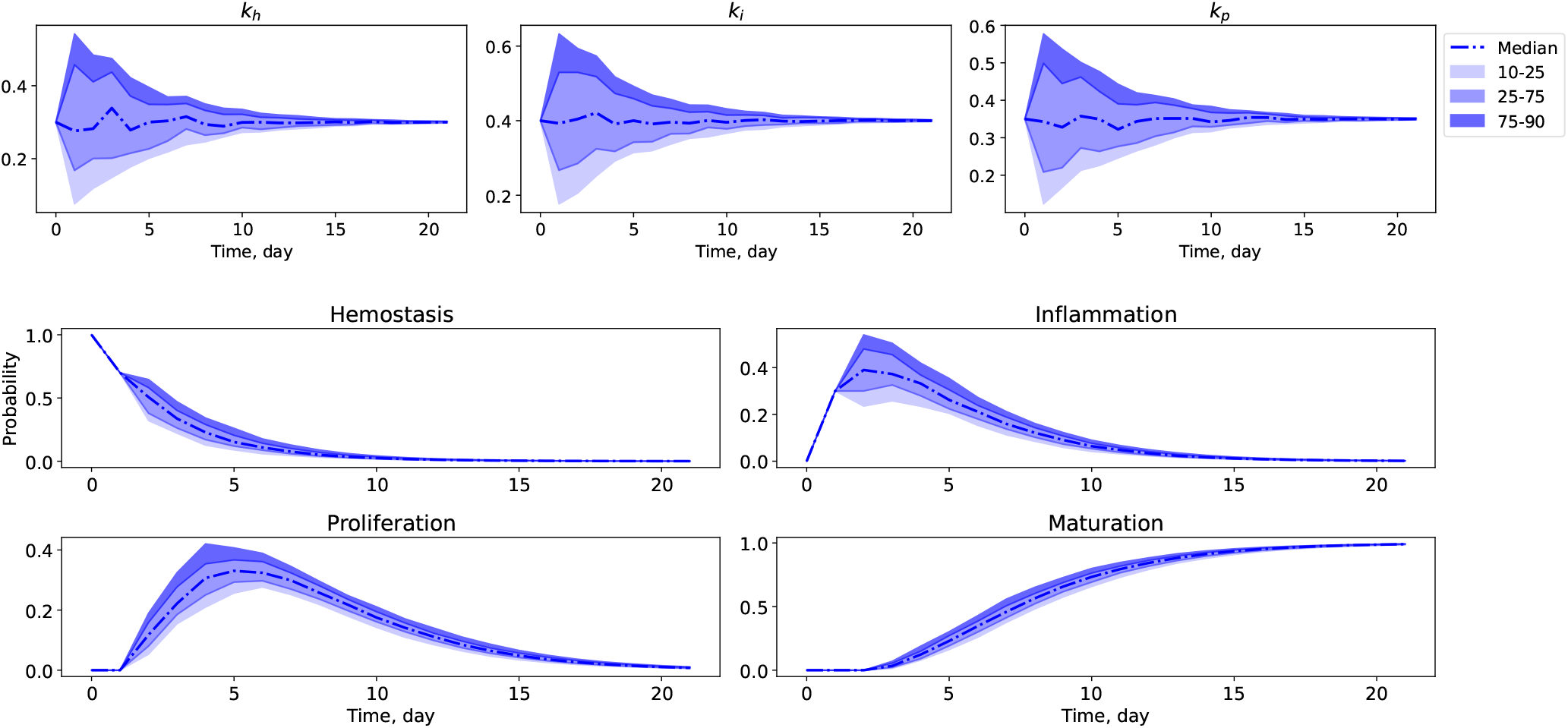
Experimental results of DeepMapper on porcine wound dataset. Values of *k*_*h*_, *k*_*i*_, *k*_*p*_ and porcine wound stage prediction across time during testing, shown by percentile.

## Discussion

The applicability of data-driven models is often constrained by the type of data they are trained on, which limits their generalization to novel or diverse datasets. For developing universal models that are applicable across different contexts, training on large and diverse datasets is essential. However, obtaining such datasets can be challenging within the scope of a single study, as data collection, labeling, and processing require substantial resources.

While it is often more practical to develop models tailored to a specific dataset or problem, this approach can be labor-intensive and time-consuming, particularly when building models from scratch. To address this challenge, we propose using a pre-trained model that can be fine-tuned and retrained for specific applications, such as clinical settings or other research projects. This transfer learning approach not only reduces the computational and data burden but also enhances the model’s applicability across different scenarios.

The proposed model’s application extends beyond determining the stages of acute wounds. We hypothesize that chronic wounds, though inherently more complex, follow a dynamic progression that may resemble the stages of acute wound healing. This suggests that identifying the stage or state of a chronic wound could inform treatment strategies, just as stage determination is critical in acute wounds. If this hypothesis holds, the model could be adapted to address a broader range of medical applications, including diagnosing and managing chronic wounds, thus extending its clinical relevance.

To further improve the model’s performance and generalizability, future work could explore the inclusion of a larger latent space and advanced techniques such as multi-head attention. These modifications would allow the model to capture more complex patterns in the data, enabling better feature representation and improving the overall accuracy in more varied wound types and conditions.

## Conclusion

In this paper, we proposed a new deep learning model, called DeepMapper, based on AutoEncoder and attention mechanism, for detecting learning a linear representation of wound healing dynamics directly from images. We demonstrate improved data efficiency in cross-species learning, enabling the model to generalize across different biological contexts with reduced data requirements. The pre-trained model, available in a public repository, offers a valuable resource for wound stage recognition in various datasets. It can serve as a foundation for developing diagnostic tools for chronic wounds, potentially improving treatment efficacy by accounting for wound stages.

In the future, we would like to explore mapping the nonlinear states into higher dimensional linear state space by considering the effects of control input through deep learning, and incorporate optimal control and reinforcement learning to derive the personalized treatment strategies to accelerate wound healing.

## ACKNOWLEDGEMENTS

This study was supported by the SciAI Center and funded by the Office of Naval Research (ONR) under Grant Number N00014-23-1-2729 and the DARPA Biotechnologies Office (DARPA/BTO) under Cooperative Agreement Number DC20AC00003.

